# SARS-CoV-2 ORF7b: is a bat virus protein homologue a major cause of COVID-19 symptoms?

**DOI:** 10.1101/2021.02.05.428650

**Authors:** Marie-Laure Fogeron, Roland Montserret, Johannes Zehnder, Minh-Ha Nguyen, Marie Dujardin, Louis Brigandat, Laura Cole, Marti Ninot-Pedrosa, Lauriane Lecoq, Beat H Meier, Anja Böckmann

**Affiliations:** Molecular Microbiology and Structural Biochemistry (MMSB), UMR 5086 CNRS/Université de Lyon, 69367 Lyon, France; Physical Chemistry, ETH Zurich, 8093 Zurich, Switzerland

## Abstract

ORF7b is an accessory protein of SARS-CoV-2, the virus behind the COVID-19 pandemic. Using cell-free synthesized ORF7b, we experimentally show that ORF7b assembles into stable multimers. The ORF7b sequence shows a transmembrane segment, which multimerizes through a leucine zipper. We hypothesize that ORF7b has the potential to interfere with important cellular processes that involve leucine-zipper formation, and present two particularly striking examples. First, leucine zippers are central in heart rhythm regulation through multimerization of phospholamban in cardiomyocytes. Second, epithelial cell-cell adhesion relies on E-cadherins, which dimerize using a transmembrane leucine zipper. Most common symptoms of SARS-CoV-2 infection, including heart arrythmias, odor loss, impaired oxygen uptake and intestinal problems, up to multiorgan failure, can be rationalized by a possible interference of ORF7b with the functions of these proteins. We ask whether this is pure coincidence, or whether our observations point to disruption by ORF7b of vital processes in COVID-19.

## Introduction

SARS-CoV-2 is a coronavirus which is at the origin of the COVID-19 pandemic. *Coronaviridae* are enveloped, single-stranded RNA viruses that infect birds and mammals, including humans. The 2019/2020 outbreak has sparked a global health crisis involving significantly more casualties than the historic severe MERS-CoV and SARS-CoV outbreaks, while other strains are relatively harmless and linked *e.g*. to the common cold. The dangerous stands were linked to recent transmission of the virus from an animal reservoir, for SARS-CoV-2 possibly from bat (*1*) *via* other mammalian hosts, to humans (*2*). The coronavirus positive-sense, single-stranded RNA (ssRNA) genome is relatively large and contains 14 open reading frames, predicted to encode 27 proteins, including four structural and eight accessory proteins (*3*). As their name suggests, accessory proteins are dispensable for viral replication, but may importantly mediate the host response to the virus, which can affect pathogenicity and virulence (*4*), e.g. through interactions with the host proteins (*5*). Accessory proteins may also be part of the viral particle (*6, 7*). It is these accessory proteins which vary greatly between variants. While structural and non-structural proteins between SARS-CoV-2 vs. SARS-CoV and SARS-CoV-2 vs. and bat-SL-CoVZXC21 are conserved over 90%, conservation of most accessory proteins falls below 90 %, and can be as low as low as 32% (*8*). One can note that the virulent strains MERS-CoV, SARS-CoV and SARS-CoV-2 have a significant number of these proteins, while there are less in more harmless coronaviruses. This points to accessory proteins as major culprits able to induce complications not observed in less virulent coronavirus infections (*4*).

ORF7b is an accessory protein of SARS-CoV-2. It shows over 93 % sequence homology with a bat coronavirus 7b protein (*5*) (Figure S1). It is a peptide composed of 43 residues, and its sequence is conserved to around 60 % between SARS-CoV and SARS-CoV-2. The ORF7b sequence clearly shows a long hydrophobic stretch which has been identified early-on as a transmembrane domain (*5*), with a luminal N-terminal and cytoplasmic C-terminal. The transmembrane domain of ORF7b has been shown to be necessary for its retention in the Golgi complex (*9*). SARS-CoV ORF7b has also been shown to be integrated in the viral particles, making it a structural component of the SARS-CoV virion (*10*). Interestingly, ORF7b has been shown to exhibit relatively high translation in cells (*11*). One can mention that 12 C-terminal residues of ORF7b are partly replaced in a SARS-CoV-2 deletion variant by 5 residues remaining from ORF8 (*12*), leaving the transmembrane domain intact.

We have recently reported ORF7b expression by wheat-germ cell-free protein synthesis, as well as NMR sample preparation (Altincekic et al., Frontiers Mol. Biosc., 2021, accepted). First solution-NMR spectra of ORF7b showed however few signals and broad resonance lines, pointing to larger protein assemblies, and indicating that membrane reconstitution and solid-state NMR approaches were needed for structural studies. We here report now that, during preliminary tests we did in the context of membrane reconstitution, ORF7b forms highly stable multimers on detergent removal, and identified a leucine zipper sequence as the likely origin. This finding places ORF7b as hypothetical interferent with cellular processes which use leucine zippers motifs in transmembrane multimerization domains. To illustrate this, we highlight two prominent cellular processes, related to heart rhythm and epithelial damage, which might be disrupted by ORF7b interfering with cellular protein interactions.

## Results

### ORF7b forms higher multimers on detergent removal

We have recently described wheat-germ cell-free protein synthesis of virtually all accessory ORFs from SARS-CoV-2 (Altincekic et al., Frontiers Mol. Biosc., 2021, accepted). ORF7b was synthesized with good yield, but was found in the pellet in absence of detergent (Figure 1a top panel). When detergents were added to the reaction, the protein was detected in the soluble fraction for all three detergents tested, namely MNG-3, Sapnov LS, and Brij-58 (Figure 1a lower panel). The protein carried a Strep-tag, and attached to Strep-Tactin magnetic beads under all conditions (SN-beads, Figure 1a). The protein can be purified in presence of DDM (Figure 1b) with a yield of 0.27 mg per milliliter wheat-germ extract used.

**Figure 1.**
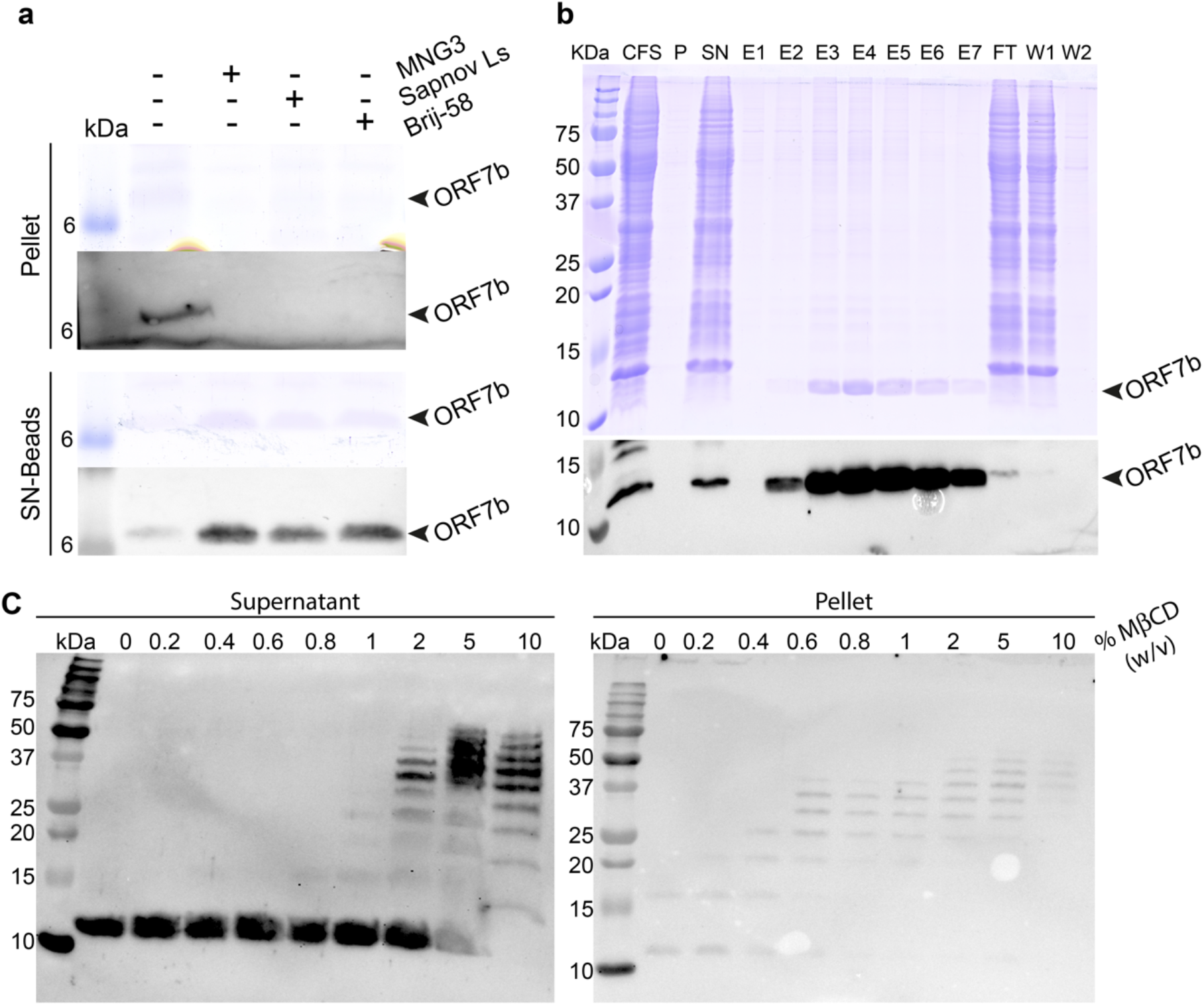
ORF7b is a membrane protein and forms multimeric assemblies. a) Wheat-germ cell-free protein synthesis of ORF7b in the absence and presence of different detergents. Pellet, pellet obtained after centrifugation of the total cell-free reaction; SN-beads, supernatant obtained after centrifugation of the total cell-free reaction and incubated with Strep-Tactin magnetic beads to capture tagged ORF7b. Comparable amounts were loaded on the gel for Pellet and SN-beads. The upper row shows Coomassie staining, the lower shows the corresponding Western blots using an antibody against the Strep-tag. Bands corresponding to ORF7b are indicated by black arrowheads. **b**) Purification of ORF7b in presence of 0.1 % DDM, Coomassie staining, and Western blots of all fractions: CFS, cell-free system; P, pellet; SN, supernatant; E, elution; FT, flow-through; W, wash. **c**) Precipitation test to determine which concentration of cyclodextrin will be needed to insert the protein into lipids. Each protein sample is subjected to increasing quantities of methyl-β-cyclodextrin (MβCD). No protein precipitates are observed in the pellet, as observed in other cases (*13*–*15*); instead, the protein oligomerizes and remains in the supernatant.

In the context of preparing ORF7b samples for solid-state NMR structural studies, the possibility to insert the protein into membranes was next explored. We use in general cyclodextrin to remove the detergent, after addition of detergent-solubilized lipids (*13*–*15*). A first step in this context is to carry out a precipitation test, to determine the amount of cyclodextrin needed to fully remove detergent from the protein. This test results generally in protein precipitates when detergent starts to be depleted from the protein (*13*–*15*). For ORF7b, however, detergent removal surprisingly, but clearly, resulted in the formation of multimers, up to approximately decamers, as observed in Western blots (Figure 1c) for around 2 % of cyclodextrin or more. The protein remained, for all cyclodextrin concentrations, in the soluble fraction, as seen in the only weak bands in the pellet fraction in Figure 1c, right panel.

These experiments clearly show that ORF7b auto assembles to form multimers, which remain intact under the denaturing conditions of the SDS gel, pointing to a very high stability.

### ORF7b shows a leucine zipper as multimerization domain

Higher-order multimers of peptides of the length of ORF7b are typically reported for viral channels, as for example the HCV p7 viroporin (*16*), the E protein from coronaviruses (*17*–*19*), and Influenza M2 (*20*). Stable multimers for some of these proteins have been detected by SDS-PAGE or Western blot, after crosslinking (*21*), and also for mutant forms (*22*). In human cells, a particularly prominent example of a multimeric transmembrane protein which conserves its multimeric form in SDS-PAGE is phospholamban (PLN), as has been shown in a mutational study (*23*) which allowed to clearly determine residues involved in the multimerization interface of the coiled coil (*24*) through the observation of pentamer bands, abolished by single alanine substitutions of leucine zipper residues.

This made us analyze in further detail OFR7b’s amino-acid sequence features, and compare it not only to different channels, but importantly to PLN, and also further regulatory peptides as sarcolipin (SLN) and myoregulin (MLN). A first analysis highlighted the exclusively hydrophobic nature of the ORF7b membrane-spanning helix, which is in contrast to the amino-acid sequences for example of Influenza M2 mentioned above, which also shows polar and charged residues in the transmembrane part. Another striking feature of the ORF7b sequence is its high abundance in Leu residues. Remarkably, exclusively hydrophobic interaction motifs involving typically Leu and Ile residues occur in the important class of leucine zippers (*25*–*27*). We manually searched ORF7b for the presence of Leu heptad repeats as typical signature of Leu zippers (*28*). And indeed, a clear succession of 4 Leu at the “a” positions, each spaced by 6 other amino acids at the “b”-”g” positions can be identified within the predicted transmembrane region of ORF7b. Interspersed with these “a” sites are on the “d” sites, important for interdigitation, I, L and V residues. While valine is not the canonical zipper motif, where dominantly leucines or isoleucines are interspersed on “d” sites, the hydrophobic Val can promote hydrophobic interactions as well.

Figure 2a shows the sequences of the analyzed peptides aligned on their respective leucine zipper region with “a” and “d” sites highlighted in red and blue respectively. The zipper sequences are plotted in Figures 2b-e as helical wheel representations. The lower panels show the multimerization via the “a” and “d” sites in a hypothetical pentamer (with the exception of PLN, for which experimental structural data is available (*29*)). One can see there how “a” and “d” amino acids form the intermolecular interactions sites.

**Figure 2:**
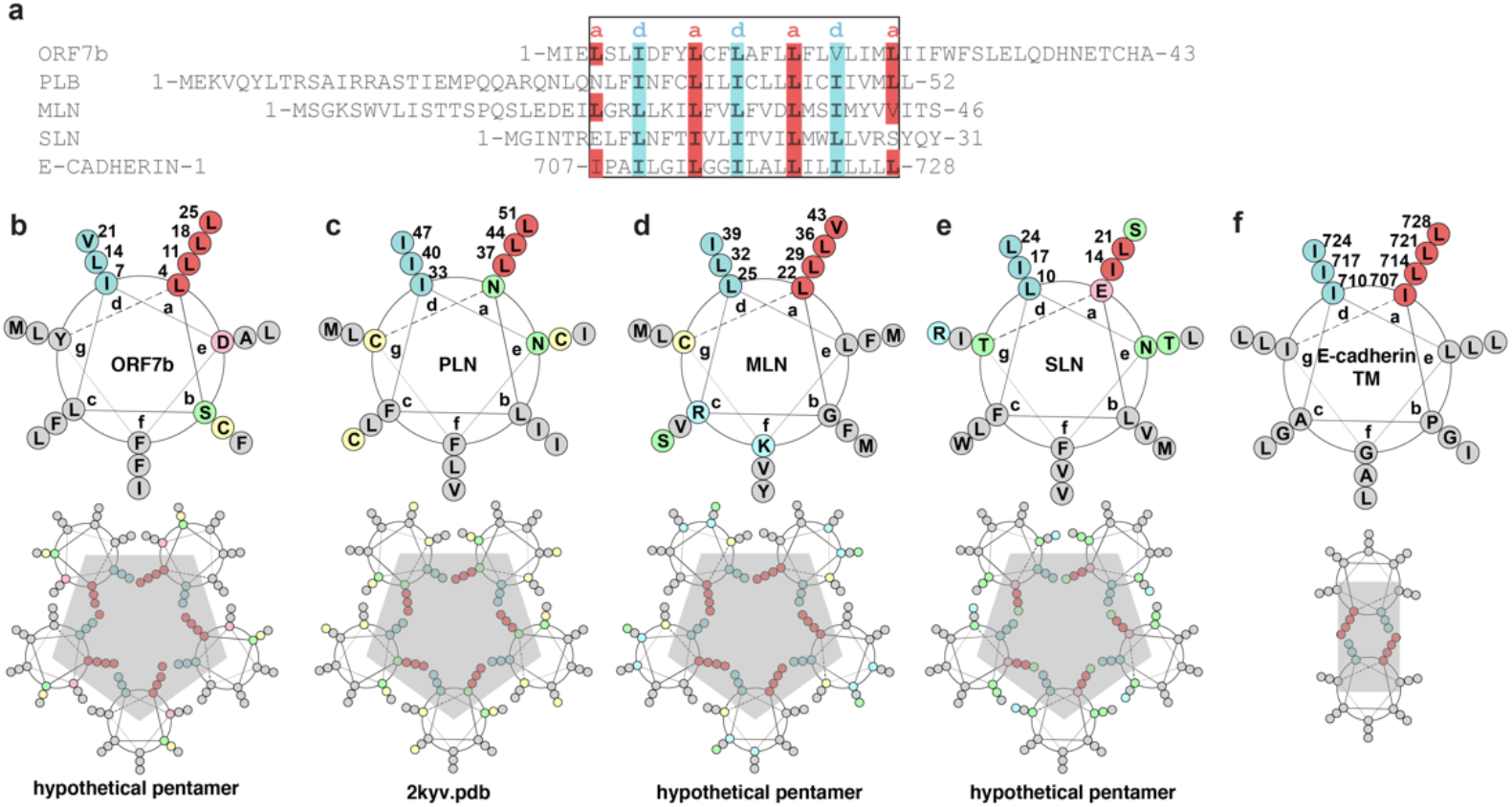
a) Sequence alignment of ORF7b, PLN, MLN, SLN and E-cadherin TM. “a” and “d” positions of the helices, forming the interfacial leucine zipper, are highlighted in red and cyan, respectively. b) Helical wheels of the five proteins, with hydrophobic positions in grey, with the exception of “a” and “d” positions which are highlighted in red and blue respectively. Polar residues are shown in green, acidic in pink, Cys in yellow, and basic in cyan. Wheels were made based on DrawCoil output (https://grigoryanlab.org/drawcoil/). While the structure of PLN is experimental (*29, 32*), the pentameric structures for ORF7, MLN and SLN are modeled based on PLN. A dimer structural model is shown for the E-cadherin TMs.

We searched for other cellular proteins where leucine zippers are of importance, and learnt that leucine zipper motifs also occur in membrane glycoproteins where they form part of the motifs driving dimerization (*30*). One well-studied example of interest in the current context is the transmembrane zipper domain of E-cadherins (*31*). E-cadherins are proteins in the epithelium which mediate cell–cell contacts, crucial namely for adhesion. The intensity of the adhesiveness is a function of the expression level of the E-cadherins, their interaction with the cytoskeleton, and their lateral interaction on the cell surface. The transmembrane domain was shown to affect E-cadherin clustering (*31*). Figure 2a shows the amino-acid sequence of the transmembrane helix of E-cadherin with “a” and “d” sites highlighted. The zipper motif is canonical in this protein, and Figure 2f shows how Leu and Ile residues can interdigitate to form the transmembrane segment dimerization.

While we extensively searched the literature for other examples, it seems that the not many more transmembrane Leu zippers have been identified so far, in contrast to their relative abundance in soluble and structural proteins (*27*). Whether this is due to them being reserved as motifs for few specialized cellular functions, or if this reflects the difficulty in searching for them especially in the transmembrane parts of sequences in large databases remained unclear.

### 3D structural ORF7b model derived from canonical leucine zipper interactions

The schematic shown in Figure 2c is perfectly verified for PLN by the published 3D NMR structure (*29*), which transmembrane part carrying the Leu zipper domain is shown in Figure 3a-c. Figure 3a shows the arrangement of the PLN transmembrane helices in the pentamer (PDB 2KYV (*29*)). Figure 3b shows how the “a” and “d” sites localize at the inner core, while the amino acids of the other five sites point into the membrane. Figure 3c shows how the “a” and “d” sites interdigitate to form the inner core of the coiled-coil pentamers, with the sites marked in the same colors as in Figure 2. The leucine and isoleucine residues tightly fill the bore of the pentamer, and the hydrophobic interactions stabilize the pentamer.

**Figure 3:**
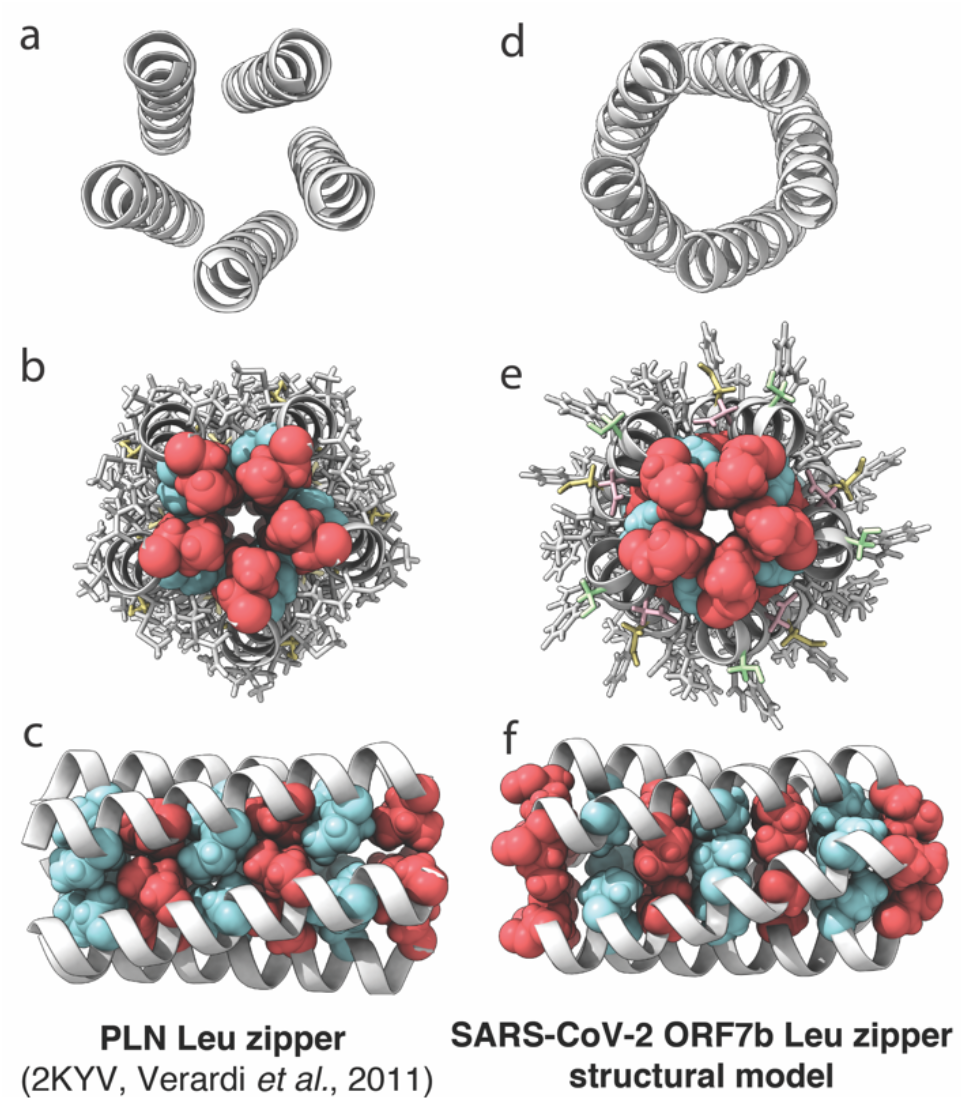
Phospholamban 3D structure (PDB 2KYV (*29*)) *versus* ORF7b transmembrane leucine zipper structural model. Shown is in panels a-c the zipper portion of the transmembrane domain of the PLN pentamer structure (PDB 2kyv (*29*)); and in d-f) the equivalent part of the ORF7b model (this work). Panels a) and d) give the top view on the coiled coil pentamer, b) and e) the same view with the zipper residues in CPK and the other residues in ball and stick. Panels c) and f) show the zipper from the side, with “a” and “d” sites highlighted in red and blue respectively (see also Fig.2). The figure was produced using ChimeraX (*34*).

Assuming that ORF7b forms indeed a leucine zipper, and thus an orientation of residues “a” and “d” in the multimer towards the inner bore similar to PLN, allowed us to build a structural model for the example of a pentameric ORF7b, which is shown in Figure 3d-f. The model was produced by the CYANA software (version 3.98.13) (*33*) in homology to the PLN structure (*29*) shown in Figure 3a-c. The α-helical secondary structure was enforced by backbone torsion-angle restraints and intramolecular distance restraints (Table S1). The 3D super structure was restrained by defined intermolecular distances between the same residue in different monomers (Table S2). Figure 3d shows the pentameric ORF7b structural model without side chains, in order to highlight the coiled-coil backbone. The orientation of the 18 ORF7b C-terminal residues cannot be modeled based on the information available. This part of the protein might form a amphipathic *α*-helix as observed for the N-terminal domain of PLN; but it might also form other structures. Figures 3e and f show that a similar arrangement of the leucine zipper in PLN is possible for the ORF7b helical bundle.

## Discussion

We here showed experimentally that ORF7b forms a multimeric structure in detergent, composed by 5-10 monomers. As the position of Leu and Ile residues in the transmembrane amino-acid sequence are highly suggestive of a leucine zipper, we propose that multimerization takes place via its transmembrane-spanning helix. Based on this assumption, we build a structural model for the ORF7b transmembrane coiled coil.

In leucine zippers, the hydrophobic leucine sidechains of two or more monomers interdigitate like the teeth of a zipper (*27*). This simple interaction motif makes them one of the most ubiquitous mediators of protein-protein interactions, which is used as a structural element to control multimerization in many cellular proteins, including many DNA interacting and also extracellular matrix and cytoskeletal networks (see Table 2 in reference (*35*)). However, transmembrane leucine-zipper interactions seem to be less abundant. They have been identified mainly in signaling and regulatory proteins, as PLN, SLN, MNL, but supposedly also DWarf Open Reading Frame (DWORF) (*36*), endoregulin (ENL) (*37*), and another-regulin (ALN) (*37*). Interestingly, they are also present in membrane glycoproteins, typically in E-cadherin, where they mediate dimer formation of the transmembrane helices. E-cadherin dimer formation is central in cell-cell adhesion.

We here emit the hypothesis that the ORF7b leucine zipper motif has the potential to interfere with the function of cellular proteins which use similar motifs. To underpin our hypothesis, we discuss two examples possibly related to COVID-19 symptoms: heart rhythm regulation by PLN and related peptides; and also E-cadherin-mediated cell-cell adhesion, at the example of interactions between sustentacular cells and olfactory neuron cilia important in the olfactory epithelium.

The finding that one of the few multimeric transmembrane Leu zippers in human cells identified today, and thus possibly a structural homologue to ORF7b, is involved in heart rate regulation intrigued us. It is known that the heart is the second target organ of the SARS-CoV-2 virus (*38, 39*), likely since it shows abundance of ACE type 2 (ACE-2) receptors which allow the virus to get internalized into the cells. Heart conditions related to SARS-CoV-2 are numerous and can arise through a variety of underlying putative mechanisms which have been put forward in the context of COVID-19 (*39*); a prominent cause includes diastolic dysfunction, which can occur due to impaired calcium transport. Defects in this activity are associated with cardiomyopathies and heart failure (*36*). Calcium traffic through sarco-endoplasmic reticulum Ca^2+^ adenosine tri-phosphatase (SERCA) is regulated by PLN, and as well by at least one other small transmembrane protein, SLN, MLN, and DWORF. The different peptides share few primary structural features, but all are predicted or determined to contain transmembrane α-helical leucine zippers. That leucine zipper domains display highly similar interaction interfaces amongst one another (Figure 2) makes it likely that they can engage in heterologous interactions as well, moreover so because the remaining residues are mainly hydrophobic in nature due to their membrane environment. Also, it has been shown for PLN that the N-terminal domains, located on the surface of the membrane, do not interact with each other (*29*), so that no specificity for homologous interactions arise from these parts neither. This leads to suppose that ORF7b can interact with, and thus possibly sequester *via* its transmembrane leucine zipper, the regulatory peptides of the PLN family (Figure 4a). It seems however unlikely that ORF7b will directly interact with SERCA, since this interaction does not occur via the Leu zipper residues, but residues outside the bore of the pentameric pore. The important functions carried out in the cell by PLN and related peptides certainly merit to be protected in organisms from interference by other proteins. A foreign protein sharing an important feature with signaling peptides like PLN might have the potential to interfere with their function.

**Figure 4.**
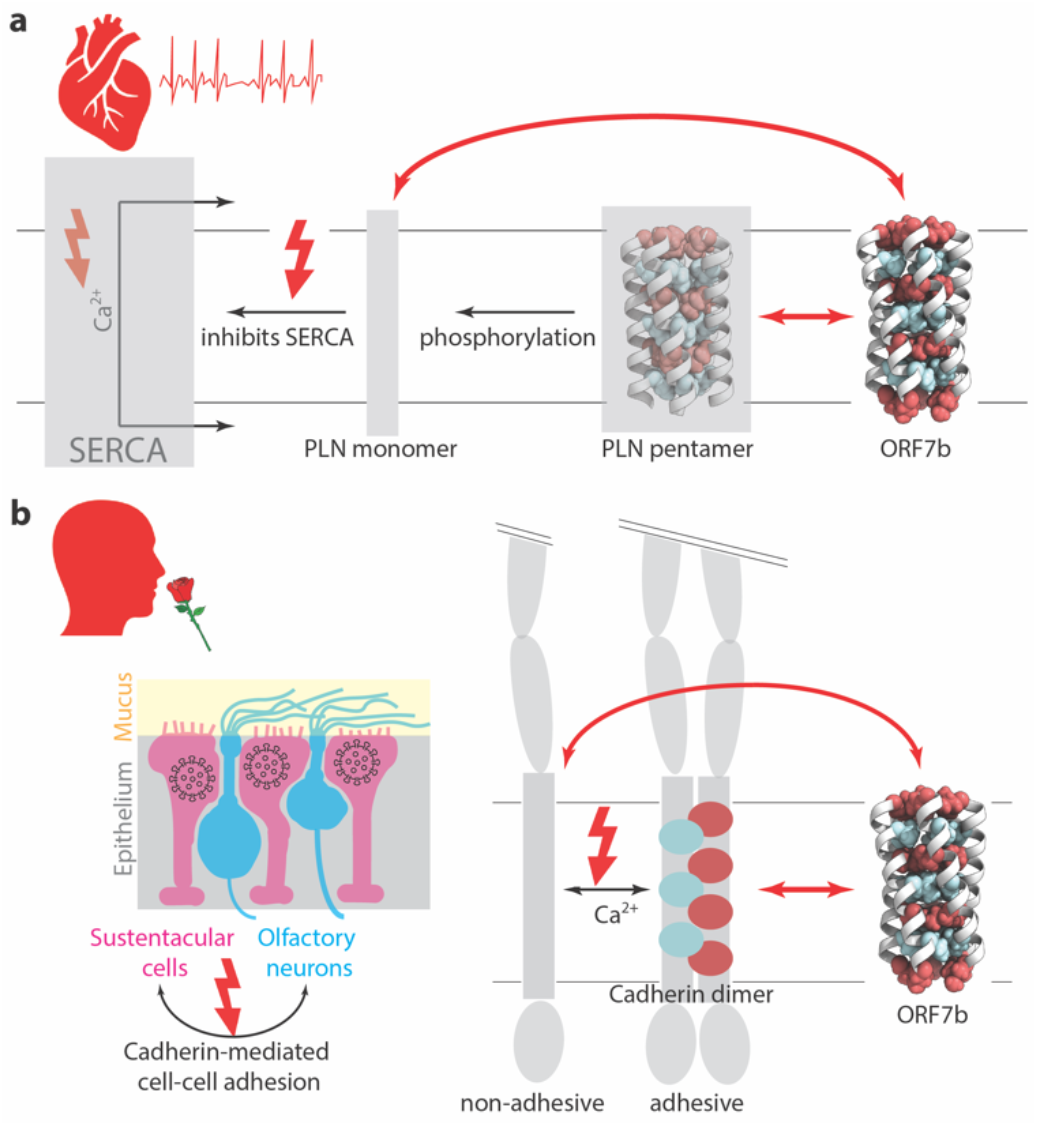
ORF7b has the potential to interfere with cellular functions. a) In the heart muscle, SERCA catalyzes Ca^2+^ transport across the membrane. PLN is an inhibitor of SERCA, and thus regulates Ca^2+^ transport, and ultimately heart muscle contraction and heart rhythm. PLN binds to SERCA as a monomer, that is formed in a phosphorylation-dependent manner. ORF7b has the potential to interact, via its leucine zipper, with PLN monomers. **b**) Olfactory neurons present cilia to the mucus. They are supported by sustentacular cells in the epithelium. Only these support cells can be entered by the virus. Cell-cell adhesion between same and different cell types is mediated by E-cadherin. For E-cadherin to be adhesive, it must dimerize, including via the leucine zipper transmembrane domain. ORF7b has the potential to interefere with adhesion by interaction of its leucine zipper with the one from the E-cadherin transmembrane domain. Cartoon of olfactory endothelium adapted from (*56*).

Another common symptom of Covid-19 is the loss of smell (*40, 41*). In a simplified picture, the olfactory epithelium is composed by support cells and sensory cells, and it is maintained by adherens and tight junctions between these cells (Figure 4b). While sensory cells have no ACE2 receptors and are likely protected from cell-entry by the virus (*40*), the virus can infect, via ACE2, support (sustentacular) cells, which present the cilia of the sensory cells (*42*). The cilia are the parts that detect odors. It has been reported recently in golden Syrian hamsters that instillation of SARS-CoV-2 in the nasal cavity resulted in transient destruction of the olfactory epithelium (*43*). It was however the support cells that were affected, not the sensory cells. A desquamation (off-peeling) of the olfactory epithelium was observed on infection, and the cilia were lost (*43*). Desquamation is due to interruption of the adhesive properties of a cell. E-cadherins present a major class of cell-cell adhesion proteins, and they must form stable dimers to be able to fulfill their adhesive function (*44*). As mentioned above, they intriguingly contain a transmembrane domain which shows a canonical leucine zipper motif (Figure 2f). The cadherin-mediated adhesive interactions between cells depend on the expression levels of the protein, the proper association of the cytosolic and cell-surface domains, as well as the transmembrane domain which was reported to affect the lateral interactions, likely *via* association of the transmembrane domains (*31*). We thus understand, with our admittedly limited insight in the topic, that disruption of this interaction by a foreign protein presenting similar interaction interfaces could perturb correct cell-cell adhesion, and result in the observed desquamation of the olfactory epithelium.

In addition to the olfactory epithelium, which damages are highly prominent in COVID-19, but far from lethal, E-cadherin is a central player in other epithelia, notably the lung and the intestine, where it equally mediates cell-cell adhesion. Disruption of this function is of importance in two debilitating medical conditions, asthma and inflammatory bowel diseases, both of which are tightly connected to E-cadherin (dys)function (*45*–*48*). Asthma results in lack of oxygen; inflammatory bowel disease results in intestinal malfunction; both are common symptoms observed in COVID-19. While the causes behind E-cadherin dysfunction in asthma / inflammatory bowel diseases are likely totally different when compared to COVID-19, the symptoms are intriguingly similar, and might both be caused by E-cadherin malfunction, as was indeed observed in SARS-CoV-2 induced intestinal responses (*49*). We thus suspect that ORF7b is perturbing E-cadherin functions, and thus is the culprit that causes the disorder in organ epithelial tissues, as for example described in (*50, 51*).

Our intention here is not to state that our possibly oversimplified view on these complex cellular processes, and the hypothetical interference caused by ORF7b, represent a fact. We rather want to report on our observations, and point out the intriguing -yet hypothetical -interactions we identified when studying ORF7b with biochemical methods. Another compellent fact is that ORF7b was not considered in latest protein interaction analyses (*52*), since its affinity purification revealed in one study an “unusually high number of background proteins and was therefore excluded” (*53*). Both E-cadherin and PLN is expressed in the HEK293 cells used in this study, and the -hypothetical -strong interaction of OFR7b with these proteins, resulting in highly stable multimers, could have caused this observation.

Our work does not yet reveal any of the functions ORF7b might take in viral fitness. It might be that ORF7b, in homology to other hydrophobic transmembrane multimers as HIV vPu or CoV E, could act as an ion channel. In any case, viroporins have been therapeutic targets in several diseases, and the druggability of these assemblies has been demonstrated in Influenza M2 (*54*) and HCV p7 (*55*).

## Conclusion

We demonstrated that ORF7b forms robust multimers, most probably stabilized by a leucine zipper. We hypothesize that this strong interaction motif enables this viral protein to interact with cellular transmembrane leucine zipper proteins, making ORF7b a major suspect for interference with cellular functions commonly observed on SARS-CoV-2 infection. More precisely, we propose that ORF7b is a suspect as causative agent for heart arrythmia and loss of smell, and advance possible molecular bases. We hypothesize that other epithelial disorders, following similar principles as odor loss, and inducing lung or bowel complications, might be caused by ORF7b as well. A drug acting on ORF7b, by stabilizing its self-interactions, or destabilizing its interactions with cellular proteins, might present an important weapon in the fight against COVID-19.

## Materials and Methods

### Plasmid

The sequence of full-length ORF7b from SARS-CoV-2 (GeneBank Accession Number 43740574) was synthesized and cloned into the pEU-E01-MCS vector (CellFree Sciences, Japan) with a SA linker and a *Strep*-tag II (SAWSHPQFEK). The resulting plasmid was amplified in *Escherichia coli* TOP10 chemically competent cells (Life Technologies). DNA was isolated using a NucleoBond Xtra Maxi kit (Macherey-Nagel, France), and was further purified by a phenol/chloroform extraction.

### Wheat germ cell-free protein synthesis

Cell-free protein synthesis was performed with a home-made wheat germ extract, as described previously (*57*–*59*). Briefly, transcription and translation steps were carried out separately. Transcription was performed for 6 hours at 37°C in Transcription buffer containing RNAsin (1U/μL), SP6 polymerase (1U/μL) (CellFree Sciences), 10 mM rNTP mix (Promega) and 0.1 μg/μL of plasmid DNA. Translation was performed for 16 hours at 22 °C without shaking using the bilayer method. Small-scale expression tests were performed in 96-well plates, while larger protein productions for further purification and lipid reconstitution were performed in 6-well plates. In the latter format, translation reaction consisted of 0.5 mL of translation mix in the bottom layer, overlaid by 5.5 mL of feeding buffer in one well. The translation mix contained 0.25 mL of transcription reaction, 0.25 mL wheat germ extract, 40 ng/mL creatine kinase, 6 mM amino acid mixture (average concentration of each amino acid 0.3 mM) and 0.1% (w/v) MNG-3 (Maltose Neopentyl Glycol-3, Anatrace). The feeding buffer contained 1x SUB-AMIX buffer (CellFree Sciences Japan), 0.1% MNG-3 and 6 mM amino acid mixture.

### SDS-PAGE and Western blotting analysis

SDS-PAGE and Western blotting analysis was performed as described in (*60*). Protein detection on the blots was performed with an antibody against the *Strep*-tag II (StrepMAB-Classic, IBA Lifesciences, Germany).

### Purification of ORF7b by affinity chromatography

After cell-free protein synthesis, the total cell-free reaction (36 mL) was incubated with benzonase, and 0.25% (w/v) DDM (N-Dodecyl-beta-Maltoside) to exchange from MNG-3 to DDM detergent, for 30 min at room temperature on a rolling wheel. Total reaction was then centrifuged at 20,000 *g* and 4 °C for 30 min, and the supernatant was loaded on a 1-mL *Strep*-Tactin column (IBA Lifesciences). Purification was performed as specified by the manufacturer, all buffers containing 0.1% DDM. ORF7b was eluted in 100 mM Tris-HCl pH8, 150 mM NaCl, 1 mM EDTA, 0.1% DDM and 2.5 mM desthiobiotin.

### Precipitation test with methyl-β-cyclodextrin

A precipitation test was performed to determine the minimal amount of methyl-β-cyclodextrin necessary to remove the detergent from the purified ORF7b sample (*61*). 13.5 μL ORF7b protein 0.20 μg/μL in 0.1% (w/w) DDM was incubated with increasing concentration of methyl-β-cyclodextrin in the range from 0.1% to 1% (w/v). In addition, the same ORF7b sample was incubated with a mixture of egg yolk phosphatidylcholine and cholesterol in a ratio 70/30 (w/w) and solubilized in Triton X-100 (detergent-to-lipid ratio of 10), and with increasing quantities of methyl-β-cyclodextrin until 10% (228 nmol). Mixtures of solubilized ORF7b protein with methyl-β-cyclodextrin were incubated for 16 hours at 4°C, and centrifuged for 1 hour at 20,000 *g*. For the highest methyl-β-cyclodextrin concentrations, centrifugation was also performed at 200,000 *g*. Presence of detergent-solubilized ORF7b protein in the supernatant or detergent-free, cyclodextrin-precipitated ORF7b protein in the pellet was analyzed by SDS-PAGE followed by Western blotting.

### Model building of ORF7b

The model of ORF7b (residues 3-25) was built in homology to PLN (PDB 2KYV) (*29*) using CYANA (*33, 62*). The input for model building were the 44 (f,y) torsion angle restraints (per monomer, identical for all monomers) set to the average values of the PLN leucine zipper structure (*29*), as well as 5 C*α*-C*α* and 5 C*β*-C*β* intra-monomer distance restraints representing the i→i+7 contacts for the zipper residues 4L, 7L, 11L, 14L, 18L, 21V and 25L (Table S1) which were set to the corresponding values in the PLN structure (pdb 2KYV). 33 inter-monomer restraint between the same resides on different monomers were used to restrain the pentamer, see Table S2. C_5_ symmetry was implemented for the structural model. Flexible linkers between the monomers were inserted into the sequence for the CYANA calculation. The 10 best structures out of the structure bundle of 100 structures showed neither violations of distance or torsion-angle restraint, nor van-der Waals. The backbone RMSD of the model-bundle is 0.15 ± 0.03 Å.

## Acknowledgement

Molecular graphics and analyses performed with UCSF ChimeraX, developed by the Resource for Biocomputing, Visualization, and Informatics at the University of California, San Francisco, with support from National Institutes of Health R01-GM129325 and the Office of Cyber Infrastructure and Computational Biology, National Institute of Allergy and Infectious Diseases. The authors acknowldge funding by the CNRS, the ETH Zurich, and by a common grant to BHM and AB from the Swiss National Science Foundation SNF (grant 31CA30_196256). BHM acknowledges support from the Swiss National Science foundation (Grant number 200020_188711), and the ERC (741863 FASTER). LL acknowleges funding from the ANR/FRM (NMR-SCoV2-ORF8) and CNRS in the form of a Momentum grant.

