## Supplemental Information for "SARS-CoV-2 ORF7b: is a bat virus protein homologue a major cause of COVID-19 symptoms?"

```
TR|E0XIZ9|E0XIZ9_9BETC         MIHLTLFDFYLCVLSLLLFLVIIMLIIFCFVLELQDLNEQ------------ 40
TR|A0A166ZLE2|A0A166ZLE2_9NIDO MNELTLIDFYLCFLAFLLFLVLIMLIIFWFSLELQDIEEPCNKVFETYRSVL 52
TR|A0A0K1Z050|A0A0K1Z050_SARS  MNELTLIDFYLCFLAFLLFLVLIMLIIFWFSLELQDIEEPCNKV-------- 44
TR|A0A6B9WFL9|A0A6B9WFL9_SARS2 MIELSLIDFYLCFLAFLLFLVLIMLIIFWFSLELQDHNETCHA--------- 43
SP|Q3LZX6|NS7B_BCHK3           MNELTLIDFYLCFLAFLLFLVLIMLLIFWFSLEIQDIEEPCNKV-------- 44
                               *  .*:*:*****.*::*****:***:**  *  **:**  :*
```

**Figure S1. ORF7b sequence alignments of SARS-CoV and bat coronaviruses**. The leucine zipper segment is underlined. E0XIZ9, Bat coronavirus BM48-31/BGR/2008; A0A166ZLE2, Bat coronavirus; A0A0K1Z050, Bat SARS-like coronavirus YNLF_31C; A0A6B9WFL9, Severe acute respiratory syndrome coronavirus 2 (2019-nCoV); NS7B_BCHK3, Bat coronavirus HKU3 (BtCoV) (SARS-like coronavirus HKU3).

**Table S1:** Intramolecular distances in PLN structure (2KYL, first structure) and used upper and lower distance limit for the CYANA calculation of the ORF7b model. upl's and lol's were set to the distance from the pdb ± 0.25 Å.

| Contact in 2KYV | Distance/ Å | Corresponding contact in Orf7b | upl / Å | lol / Å |
|---|---|---|---|---|
| A 33I Cα – A 40I Cα | 10.56 | A 7I Cα – A 14L Cα | 10.81 | 10.31 |
| A 33I Cβ – A 40I Cβ | 10.65 | A 7I Cβ – A 14L Cβ | 10.90 | 10.40 |
| A 40I Cα – A 47I Cα | 10.19 | A 14L Cα – A 21V Cα | 10.44 | 9.94 |
| A 40I Cβ – A 47I Cβ | 10.14 | A 14L Cβ – A 21V Cβ | 10.39 | 9.89 |
| A 30N Cα – A 37L Cα | 10.38 | A 4L Cα – A 11L Cα | 10.63 | 10.13 |
| A 30N Cβ – A 37L Cβ | 10.35 | A 4L Cβ – A 11L Cβ | 10.60 | 10.10 |
| A 37L Cα – A 44L Cα | 10.45 | A 11L Cα – A 18L Cα | 10.70 | 10.20 |
| A 37L Cβ – A 44L Cβ | 10.44 | A 11L Cβ – A 18L Cβ | 10.69 | 10.19 |
| A 44L Cα – A 51L Cα | 10.30 | A 18L Cα – A 25L Cα | 10.55 | 10.05 |
| A 44L Cβ – A 51L Cβ | 10.57 | A 18L Cβ – A 25L Cβ | 10.82 | 10.32 |

**Table S2:** Measured intermolecular distances in PLN structure (2KYL, first structure) and used upper and lower distance limit for the CYAN calculation of the ORF7b model.

| Contact in 2KYV | Distance / Å | Corresponding contact in Orf7b | upl / Å | lol / Å |
|---|---|---|---|---|
| A 29Q Cα – B 29Q Cα | 8.59 | A 3E Cα – B 3E Cα | 8.84 | 8.34 |
| A 29Q Cα – C 29Q Cα | 13.91 | A 3E Cα – C 3E Cα | 14.16 | 13.66 |
| A 30N Cα – B 30N Cα | 7.88 | A 4L Cα – B 4L Cα | 8.13 | 7.63 |
| A 30N Cβ – B 30N Cβ | 8.38 | A 4L Cβ – B 4L Cβ | 8.63 | 8.13 |
| A 31L Cα – B 31L Cα | 11.72 | A 5S Cα – B 5S Cα | 11.97 | 11.47 |
| A 32F Cα – B 32F Cα | 11.17 | A 6L Cα – B 6L Cα | 11.42 | 10.92 |
| A 33I Cα – B 33I Cα | 7.02 | A 7L Cα – B 7L Cα | 7.27 | 6.77 |
| A 33I Cβ – B 33I Cβ | 5.60 | A 7L Cβ – B 7L Cβ | 5.85 | 5.35 |
| A 34N Cα – B 34N Cα | 9.17 | A 8D Cα – B 8D Cα | 9.42 | 8.92 |
| A 35F Cα – B 35F Cα | 12.02 | A 9F Cα – B 9F Cα | 12.27 | 11.77 |
| A 36C Cα – B 36C Cα | 9.36 | A 10Y Cα – B 10Y Cα | 9.61 | 9.11 |
| A 37L Cα – B 37L Cα | 6.81 | A 11L Cα – B 11L Cα | 7.06 | 6.56 |

| | | | | |
|---|---|---|---|---|
| A 37L Cβ – B 37L Cβ | 6.38 | A 11L Cβ – B 11L Cβ | 6.63 | 6.13 |
| A 38I Cα – B 38I Cα | 10.90 | A 12C Cα – B 12C Cα | 11.15 | 10.65 |
| A 39L Cα – B 39L Cα | 11.32 | A 13F Cα – B 13F Cα | 11.57 | 11.07 |
| A 40I Cα – B 40I Cα | 7.23 | A 14L Cα – B 14L Cα | 7.48 | 6.98 |
| A 40I Cβ – B 40I Cβ | 5.66 | A 14L Cβ – B 14L Cβ | 5.91 | 5.41 |
| A 41C Cα – B 41C Cα | 9.10 | A 15A Cα – B 15A Cα | 9.35 | 8.85 |
| A 42L Cα – B 42L Cα | 12.25 | A 16F Cα – B 16F Cα | 12.50 | 12.00 |
| A 43L Cα – B 43L Cα | 9.73 | A 17L Cα – B 17L Cα | 9.98 | 9.48 |
| A 44L Cα – B 44L Cα | 7.35 | A 18L Cα – B 18L Cα | 7.60 | 7.10 |
| A 44L Cβ – B 44L Cβ | 6.86 | A 18L Cβ – B 18L Cβ | 7.11 | 6.61 |
| A 45I Cα – B 45I Cα | 11.56 | A 19F Cα – B 19F Cα | 11.81 | 11.31 |
| A 46C Cα – B 46C Cα | 12.12 | A 20L Cα – B 20L Cα | 12.37 | 11.87 |
| A 47I Cα – B 47I Cα | 8.09 | A 21V Cα – B 21V Cα | 8.34 | 7.84 |
| A 47I Cβ – B 47I Cβ | 6.54 | A 21V Cβ – B 21V Cβ | 6.79 | 6.29 |
| A 48I Cα – B 48I Cα | 9.55 | A 22L Cα – B 22L Cα | 9.80 | 9.30 |
| A 49V Cα – B 49V Cα | 13.04 | A 23I Cα – B 23I Cα | 13.29 | 12.79 |
| A 50M Cα – B 50M Cα | 11.45 | A 24M Cα – B 24M Cα | 11.70 | 11.20 |
| A 51L Cα – B 51L Cα | 7.98 | A 25L Cα – B 25L Cα | 8.23 | 7.73 |
| A 51L Cβ – B 51L Cβ | 6.58 | A 25L Cβ – B 25L Cβ | 6.83 | 6.33 |
| A 52L Cα – B 52L Cα | 11.31 | A 26I Cα – B 26I Cα | 11.56 | 11.06 |
| A 52L Cα – C 52L Cα | 18.32 | A 26I Cα – C 26I Cα | 18.57 | 18.07 |
